# K18-hACE2 Mice for Studies of COVID-19 Treatments and Pathogenesis Including Anosmia

**DOI:** 10.1101/2020.08.07.242073

**Authors:** Jian Zheng, Lok-Yin Roy Wong, Kun Li, Abhishek K. Verma, Miguel Ortiz, Christine Wohlford-Lenane, Mariah R. Leidinger, C. Michael Knudson, David K. Meyerholz, Paul B. McCray, Stanley Perlman

## Abstract

The ongoing COVID-19 pandemic is associated with substantial morbidity and mortality. While much has been learned in the first months of the pandemic, many features of COVID-19 pathogenesis remain to be determined. For example, anosmia is a common presentation and many patients with this finding show no or only minor respiratory signs. Studies in animals experimentally infected with SARS-CoV-2, the cause of COVID-19, provide opportunities to study aspects of the disease not easily investigated in human patients. COVID-19 severity ranges from asymptomatic to lethal. Most experimental infections provide insights into mild disease. Here, using K18-hACE2 mice that we originally developed for SARS studies, we show that infection with SARS-CoV-2 causes severe disease in the lung, and in some mice, the brain. Evidence of thrombosis and vasculitis was detected in mice with severe pneumonia. Further, we show that infusion of convalescent plasma (CP) from a recovered COVID-19 patient provided protection against lethal disease. Mice developed anosmia at early times after infection. Notably, while treatment with CP prevented significant clinical disease, it did not prevent anosmia. Thus K18-hACE2 mice provide a useful model for studying the pathological underpinnings of both mild and lethal COVID-19 and for assessing therapeutic interventions.

## INTRODUCTION

The COVID-19 pandemic is associated with more than 18 million cases and more than 650,000 deaths worldwide since its emergence in December of 2019. SARS-CoV-2 uses the same receptor as SARS-CoV, human angiotensin-converting enzyme 2, (hACE2)^1^. Many of the animals infectable by SARS-CoV can also be experimentally infected with SARS-CoV-2, but these animals generally develop mild disease^2-6^. Conversely, mice, which are SARS-CoV-susceptible, are resistant to infection with SARS-CoV-2 because of incompatibilities between mouse ACE2 and the viral spike protein^7^. Several approaches have been or will be used to sensitize mice to infection, including providing hACE2 by adenovirus transduction^8,9^, mutating SARS-CoV-2 spike protein so that it binds to mACE2^10^, and modifying mACE2 so that cells are susceptible to SARS-CoV-2^11^. During the 2003-2004 SARS epidemic, since mice developed only mild disease, we and others engineered mice that transgenically displayed hACE2^12-14^. K18-hACE2 mice express hACE2 driven by cytokeratin 18 promoter, predominantly in epithelial cells^15^. SARS-CoV infected-K18-hACE2 mice developed an overwhelming encephalitis, with lung disease characterized by aspiration pneumonia and evidence of mild virus-induced pneumonia^12^. Both SARS-CoV-2 and SARS-CoV use hACE2 but show differences in disease manifestations in patients. SARS-CoV-2 infects the upper airways to a much greater extent than SARS-CoV and has been associated with clinical manifestations such as anosmia, ageusia, thrombosis and endothelial damage in the lung vasculature, cardiac and neurological disease, and a multisystem inflammatory disease in children and adolescents^16-18^. Based on these observations, we reasoned that SARS-CoV-2 infection of the K18-hACE2 mice might also show differences in pathogenesis. Here we demonstrate the utility of K18-hACE2 mice for studies of COVID-19 disease and treatments.

## RESULTS

### K18-hACE2 transgenic mice develop significant disease following SARS-CoV-2 infection

To assess the susceptibility of K18-hACE2 mice to SARS-CoV-2, we performed an intranasal challenge using inocula of 10^3^, 10^4^, and 10^5^ PFU. As shown in Fig. 1a, we observed a dose-dependent increase in weight loss and mortality. At the 10^5^ PFU inoculum, all animals succumbed, while there was variable mortality when mice were inoculated with 10^4^ PFU. To better understand the cause(s) of this morbidity and mortality, we assessed the tissue distribution of viral genomic RNA by qRT PCR at 2, 4, and 6 days post infection (dpi) in animals receiving the 10^5^ PFU inoculum. The predominant target organs were the lung at early timepoints, and variably, the brain at later timepoints (Fig. 1b, left panel). We also detected viral RNA in heart, liver, spleen, kidney, intestine, and colon. To determine which tissues were sites of virus replication, we performed plaque assays (Fig. 1b, right panel). The virus replicated to high titers in lung tissue at 2 dpi and decreased at 4 and 6 dpi. In some, but not all animals, the brain tissue titers gradually increased from 2 to 6 dpi. There was limited replication in the kidney, small intestine, and colon, perhaps signifying low level hematogenous dissemination. We note that the K18 promoter predominantly directs gene expression to multiple epithelia-lined tissues^15^. While K18 is an epithelial cell protein, K18-based expression of a LacZ reporter was also reported in cortical and brainstem neurons^19^. These results demonstrate that SARS-CoV-2 infection of K18-hACE2 mice causes dose-dependent illness associated with significant involvement of lungs with a subset of animals developing brain infection at higher inocula (10^5^ PFU).

**Fig. 1.**
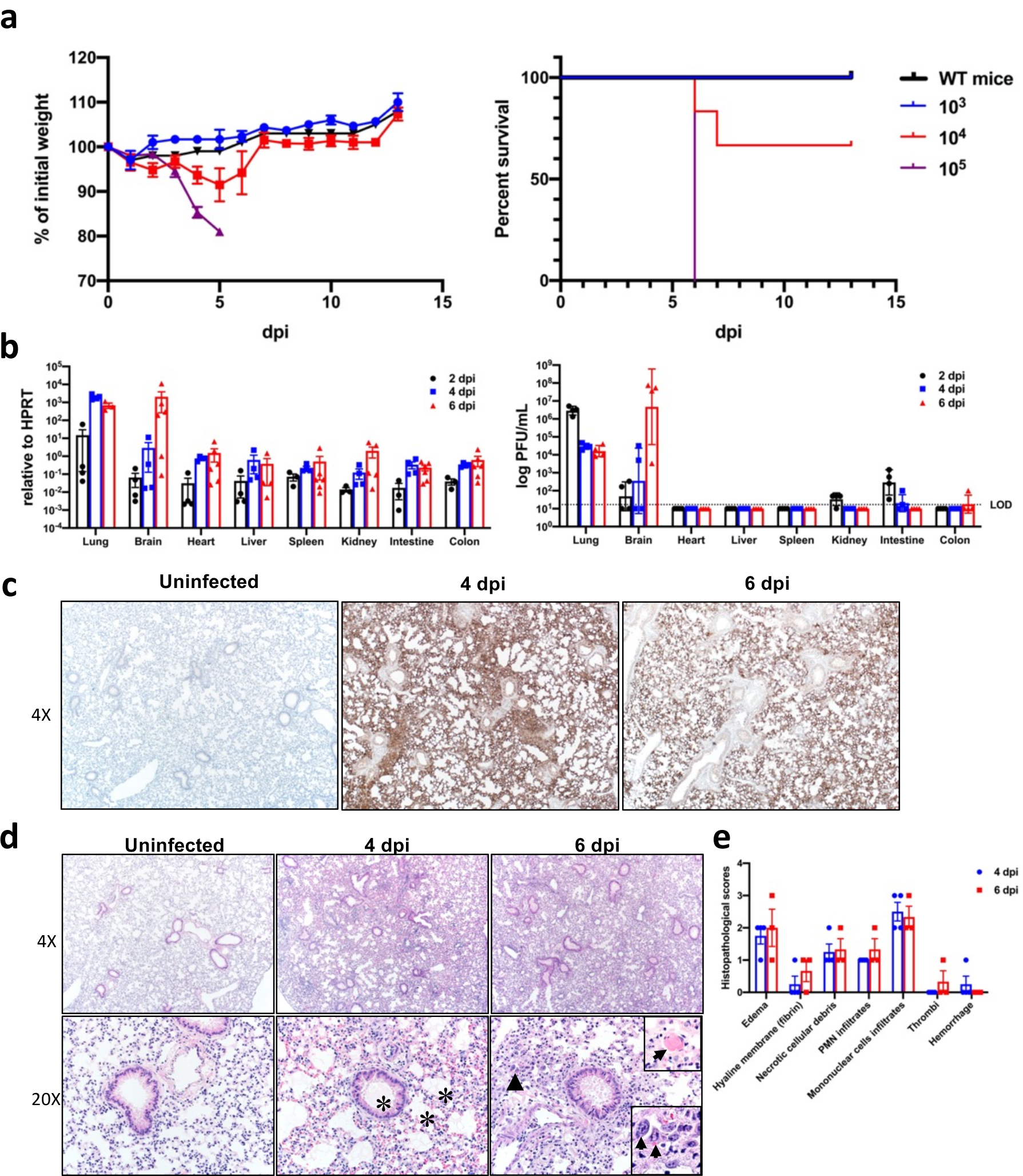
Clinical and pathological disease in K18-hACE2 transgenic mice following SARS-CoV-2 infection. a. Percentage of initial weight and survival of K18-hACE2 mice infected with 10^3^, 10^4^, and 10^5^ PFU SARS-CoV-2/mouse. b. Viral RNA detected by qPCR targeting viral N gene with normalization to HPRT for the indicated organs at 2, 4, and 6 dpi (left panel). Infectious virus titers detected by plaque assay in different organs at 2, 4, and 6 dpi (right panel). LOD=limit of detection. c. Lungs from uninfected (n=3), and infected (day 4 (n=4) and day 6 (n=3) p.i.) were analyzed by immunohistochemistry using anti-SARS-CoV-2 N antibody. d. Sections of paraffin-embedded lungs were stained with H&E. Note airway edema and alveolar hyaline membranes (asterisks, middle bottom panel), vascular thrombosis (inset, top right, 6 dpi 20x panel), dying cells with pyknotic to karyorrhectic nuclei, and a proliferative alveolar epithelium with mitotic figures (arrowhead and inset, lower right, 6 dpi 20x panel). e. Summary of histological scoring in the lungs, as described in Experimental Procedures.

### The SARS-CoV-2 pulmonary infection is extensive and progressive

In view of the early pulmonary virus replication, we stained lungs at 4 and 6 dpi for SARS-CoV-2 antigen. Using an inoculum of 10^5^ PFU, we observed extensive and diffuse parenchymal localization of SARS-CoV-2 nucleocapsid (N) protein at 4 dpi that was less intense by 6 dpi (Fig. 1c). Uninfected tissues showed no N protein signal. Examination of hematoxylin and eosin stained tissues revealed extensive evidence of diffuse alveolar damage with progressive alveolar/interstitial lesions characterized by edema, inflammation, and focal cytomegaly in some alveolar lining cells (Fig. 1d). Also observed was an accumulation of immune effector cells, including granulocytes and macrophages, evidence of cell death, hemorrhage, hyaline membranes, occasional vascular thrombi, and rare syncytia. Pulmonary vascular thrombi are often observed in severe human COVID-19^16^. Between 4 and 6 dpi there was further progression of the findings of diffuse alveolar damage. Alveolar septal thickening resulted from increased cellularity composed of infiltrating inflammatory cells, dying cells with pyknotic to karyorrhectic nuclei, and a proliferative alveolar epithelium with mitotic figures (Fig. 1d). These findings are further summarized by histopathologic scoring at 4 and 6 dpi (Fig. 1e).

We also surveyed liver, heart, spleen, kidney, small intestine, and colon tissues for disease associated changes (Extended Data Fig. 2). In the liver, we observed blood vessels filled with aggregates composed of erythrocytes and platelets. At 4 dpi, occasional vessels had evidence of fibrin thrombi adherent along the vascular wall (Extended Data Fig. 2a), suggestive of the coagulopathy observed in COVID-19 autopsies ^20^. The other tissues examined showed no disease-specific changes. To further characterize the lung tissue responses to SARS-CoV-2 infection, we surveyed the expression of several cytokines, chemokines, and innate immune transcripts at 2, 4, and 6 dpi by qRT PCR. As shown in Fig. 2a, there were significant increases in type I, II, and III interferons, ISG-15, RIG-I, MDA5, IL-1β, IL-6, IL-8, IL-12, CCL-2, CCL-5, CCR7, CXCL-2, CXCL-9, CXCL-10, and TNF compared to uninfected lungs over the course of infection.

**Fig. 2.**
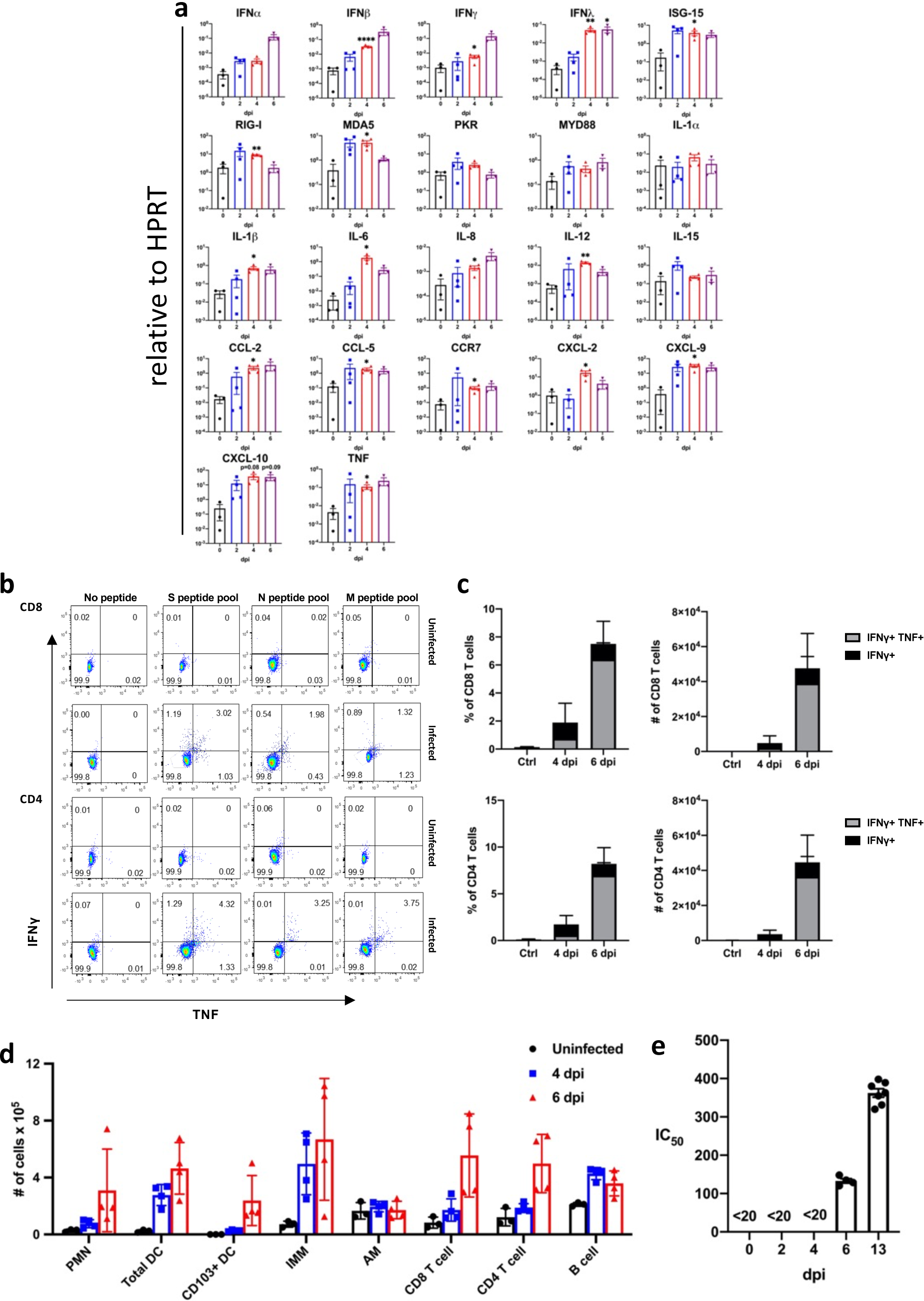
Inflammatory mediators and immune effector cells contribute to lung disease phenotype. a. Cytokine and chemokine transcripts were measured by qPCR following reverse transcription of RNA isolated from the lungs of SARS-CoV-2 infected K18-hACE2 mice (n=3 or 4 for each time point). Statistical significance compared to results obtained at 0 dpi. b. Representative FACS plot of IFNγ+TNF+ CD8 and CD4 T cells after stimulated with indicated peptide pools) in the lungs of 10^5^ PFU SARS-CoV-2 infected K18-hACE2 mice. c. Summary data are shown (n=3 mice/time point). d. Quantification of immune cells in the lungs (n=3 for uninfected group and n=4 for 4 and 6 dpi). e. Sera were collected from infected mice at the indicated time points and IC_50_ values determined by neutralization of SARS-CoV-2 pseudoviruses expressing luciferase.

### Lung disease is characterized by an influx of immune effector cells

To further investigate the factors contributing to lung injury and virus clearance, we performed immunophenotyping of infiltrating cells at 4 and 6 dpi. As shown in Fig. 2b-d, we observed a progressive increase in the numbers of macrophages and monocytes, neutrophils, and CD4 and CD8 T cells. Infiltrating inflammatory macrophage/monocytes and neutrophils may have both protective and pathogenic roles, based on previous studies of SARS and MERS^21^. Virus-specific T cells, which are required for optimal SARS-CoV-2 clearance, were detected in lungs at 6 dpi. Virus-specific CD8 and CD4 T cells responded to peptide pools representing SARS-CoV-2 S, N, and M proteins to varying extents, as assessed by expression of IFN-γ and TNF respectively (Fig. 2b, c). Together these results indicate that lung disease in K18-hACE2 mice shares features with severe COVID-19. Another feature of severe human COVID-19 is an early antibody response, consistent with plasmablast activation^22^. We detected serum neutralizing antibodies as early as 6 dpi, consistent with such a response (Fig. 2e).

### SARS-CoV-2 causes brain infection in some K18-hACE2 mice

As shown in Fig. 1b, infectious virus was detected in the brains of most mice following 10^5^ PFU intranasal inoculation. To investigate this in more detail, we immunostained tissues for the viral N protein at 4 and 6 dpi. As shown in Fig. 3a, we observed no antigen staining at 4 dpi (4/4 brains studied) but at 6 dpi saw extensive staining in several brain regions including the olfactory bulb, cerebral cortex, caudate/putamen, thalamus, hypothalamus, and ventral striatum (2/3 brains studied) (Extended Data Table 1). Uninfected animals showed no N protein staining (3/3 animals studied). Many of these areas are secondary or tertiary connections of the olfactory bulb, consistent with virus entry at this site. H&E staining demonstrated cell death and thrombi in the thalamus (Fig. 3b, c) and neutrophilic and monocytic infiltrates in the meninges as well as cells undergoing degeneration (Fig. 3d). The thalamus is affected in many COVID-19 patients with neurological disease^23^. Of note, while most of the affected areas were connected to the olfactory bulb, some, such as the area postrema and hypoglossal nucleus, are not directly connected. Similar results were observed in analyses of SARS-CoV-infected mice^24^. These results revealed an important role for olfactory bulb infection in spread to the brain in some mice but also indicated that virus may enter the central nervous system by other routes. Infection of the olfactory epithelium is expected to precede brain infection and could also contribute to the anosmia and ageusia observed in many patients, including some who are asymptomatic or mildly symptomatic.

**Fig. 3.**
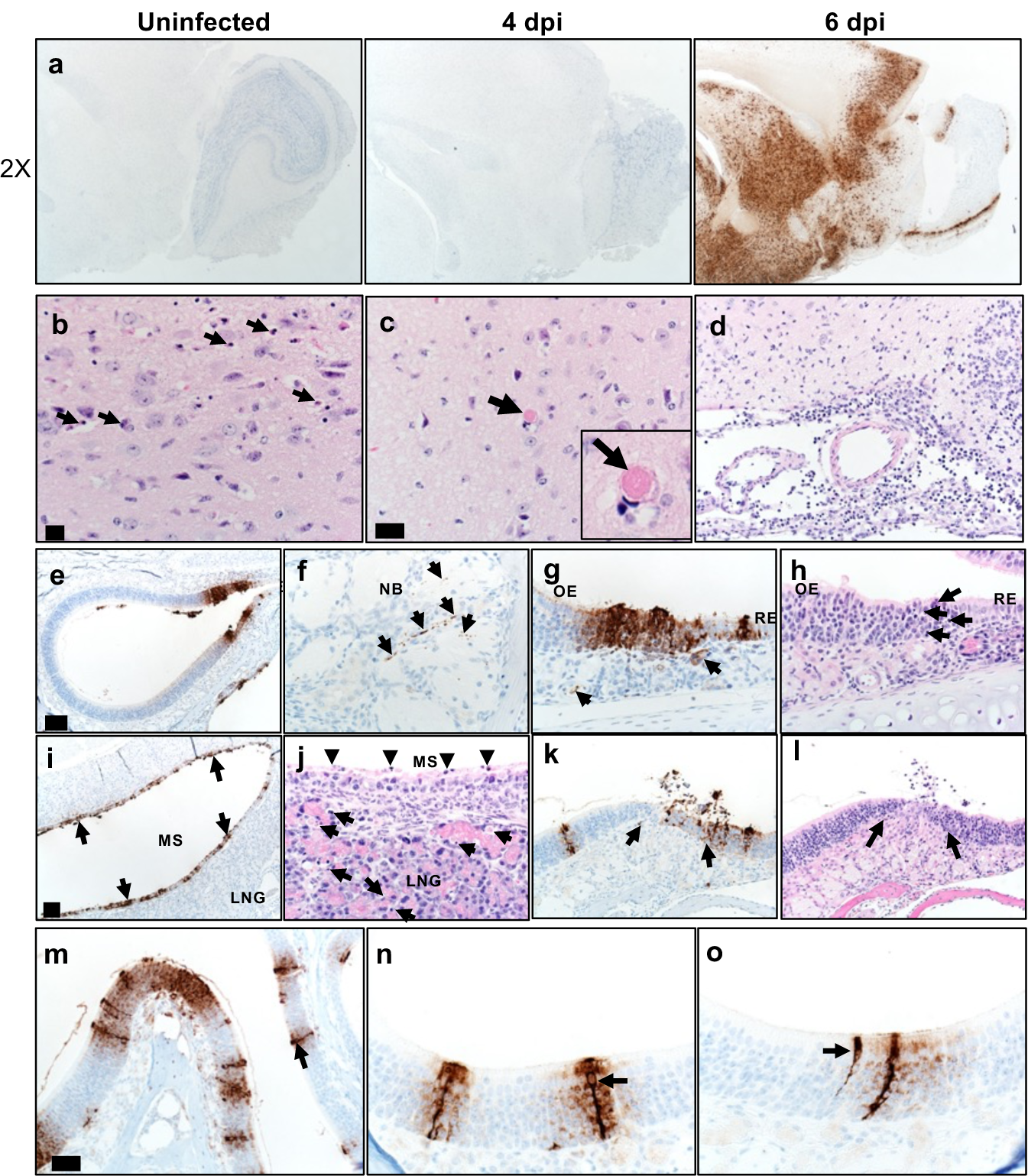
Some K18-hACE2 mice develop brain disease after SARS-CoV-2 infection. a. Brains were prepared from uninfected and infected mice and analyzed for SARS-CoV-2 by immunohistochemistry. b, c. Multiple sites of dead cells (arrows) characterized by cellular and karyorrhectic nuclear debris (b, arrows) and of thrombi (c, arrow and inset) in thalamus. Bar = 17 µm. d. Examination of meninges at day 6 p.i. revealed increased cellularity composed of degenerate cells and neutrophils and mononuclear cells. Cellular and karyorrhectic nuclei debris were also detected in the perivascular regions (200X). e-o. Nasal and sinus tissue were examined at days 2 (e-j) and 5 (k-o) p.i. e-h. Olfactory epithelium (OE) was stained with anti-SARS-CoV-2 N antibody (e-g) or H&E stain (h). f. Nerve bundles (NB) subjacent to OE had punctate immunostaining (brown, arrows). g. Subjacent to sites of immunostaining for N protein (brown, black arrows), endothelial lining of vessels (left arrow) and bowman glands (right arrow) were occasionally positive for virus antigen (brown). h. Sites of N protein localization in OE had evidenced of cell death and cellular debris (arrows). Bar = 100 µm (e) and 25 µm (f-h). i-j. Maxillary sinus (MS) stained with anti-SARS-CoV-2 N antibody (i) or H&E (j). i. MS lining epithelium had extensive immunostaining for N protein (brown, arrows). j. ME epithelium had common sloughing and cellular debris (arrowheads). The lateral nasal glands (LNG) also had multifocal cellular and karyorrhectic debris (arrows). Bar = 75 µm (i) and 19 µm (j), respectively. k-l. Olfactory epithelium still had foci of SARS-CoV-2 N protein immunostaining at d. 5 p.i. (k, arrows, left) that was often localized near interface with respiratory epithelium. In these sites, there was cellular sloughing and loss of cellularity (arrows, right). 200x. m-o. Arrows point to “classic” morphology and strong labeling of sustentacular cells with expanding labeling of adjacent cells. Bar = 50 (m) and 25 µm (n, o).

### SARS-CoV-2 productively infects the sinonasal epithelium including sustentacular cells

To assess virus replication in the upper respiratory tract following intranasal inoculation, we quantified viral RNA in nasal secretions by qRT PCR at 3 dpi. We observed that secretions from 5 of 7 mice showed evidence of SARS-CoV-2 genomic RNA (C_t_ =21.5 +/-1.2 (mean +/- SEM)), and of these 5, 4 had evidence of subgenomic RNA (36.7 +/-1.1 (mean +/- SEM), consistent with active virus replication. Viral antigen was readily detected in both the respiratory and olfactory epithelium at 2 and 5 dpi. At 2 dpi virus antigen was present at multiple sites, often at the interface of the olfactory and respiratory epithelium (Fig. 3e). Viral antigen was also detected in nerve bundles subjacent to the olfactory epithelium (Fig. 3f) and occasionally in vascular endothelia (Fig. 3g, left arrow) and Bowman’s glands (Fig. 3g, right arrow). At sites of antigen positivity in the olfactory epithelium and maxillary sinus, we observed cell death and cellular debris at day 2 (Fig. 3h-j), which progressed to cell sloughing and loss of cellularity by 5 dpi. (Fig. 3k, l). ACE2 has been detected in sustentacular cells in the olfactory epithelium^25^, but not in olfactory sensory neurons, suggesting that these cells are a primary site of infection. Consistent with this, we detected SARS-CoV-2 antigen in sustentacular cells (Fig. 3m-o). Infection of sustentacular cells is not expected to result in spread to the olfactory bulb and its connections, but could still contribute to anosmia.

### SARS-CoV-2 infected K-18-hACE2 mice exhibit anosmia

To directly assess anosmia, we performed two sets of behavioral tests, both of which require a normal sense of smell, as described in Experimental Procedures. First, male mice were exposed to bedding containing female or male dander in a 2 ml Eppendorf tube (Fig. 4a). Mice identify the tube visually, and then preferentially spend time with the female dander, if olfaction is normal. In a second experiment, we used a buried food test, in which mice are attracted to a food item hidden in the bedding that they were previously conditioned to detect (Fig. 4c). K18-hACE2 mice infected with SARS-CoV-2 did worse than controls in both tests so that at days 2 and 3 p.i., they spent less time in the vicinity of the female dander (Fig. 4b) and took longer to find buried food (Fig. 4d, e). At 2 and 3 dpi, brains were not infected (Fig. 1b). Additionally, mobility was largely normal since there were no differences in the amount of time spent in exploring the tube containing male dander, when infected or uninfected mice were compared (Fig. 4b). This suggests that anosmia at these times points is primarily caused by infection of the nasal epithelium and is not a consequence of spread to the brain.

**Fig. 4.**
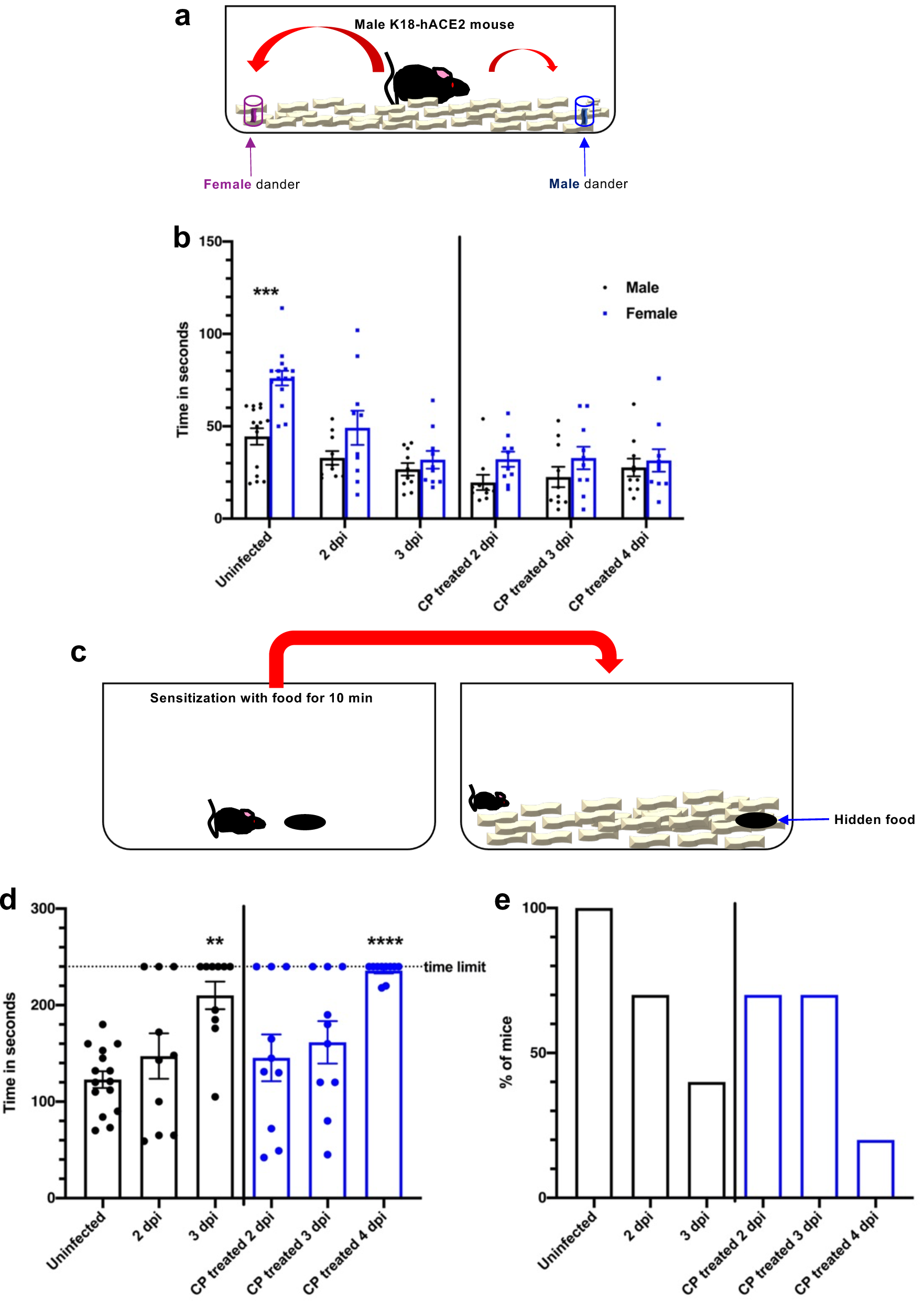
SARS-CoV-2 infection causes anosmia in K18-hACE2 mice. a. Schematic showing social scent discrimination test. b. Each male mouse was allowed 5 minutes in the cage with male and female scent (bedding from male or female cages) placed at two corners of the cage. The time that male mice spent sniffing male or female scent was recorded. Data were analyzed by 2-way ANOVA. ***p<0.001. In some experiments, mice were pretreated with undiluted convalescent plasma (results on right of vertical line). c. Schematic showing buried food test. d. Buried food test. Food was buried under the bedding and each mouse was allowed 4 minutes in the cage to search for the food. The dotted line shows the time limit of 4 minutes. Data were analyzed by 1-way ANOVA. ****p<0.0001. e. The percentage of mice that found the buried food within 4 minutes is shown. In some experiments, mice were pretreated with undiluted convalescent plasma (denoted by blue bars in d, e).

### Human COVID-19 convalescent plasma protects mice from severe clinical disease but not from anosmia

Convalescent plasma (CP) from COVID-19 survivors is being tested as a treatment in patients with clinical COVID-19^26^. Controlled studies are underway but results of these studies have not yet been released. Studies have demonstrated positive effects of CP on outcomes in several infectious diseases, if titers are sufficiently high and if CP is administered early in the disease course^27^. To assess the effect of CP on outcomes in K18-hACE2 mice infected with SARS-CoV-2, we administered undiluted and diluted human high titer plasma (neutralizing titer 1:1,480) 12 hours before infection. Undiluted plasma administered 12 hours before infection protected mice from death but not mild weight loss and reduced lung tissue titers while CP diluted 1:3 provided partial protection (Fig. 5a, c). Convalescent plasma pretreatment markedly inhibited spread of infection to the brain (Fig. 5c). On examination of lung tissues, levels of viral antigen and pathological changes were greatly decreased by CP treatment (Fig. 5d, e). Treatment did not seem to affect primary infection of lung cells but rather prevented secondary spread within the lung. As shown in Fig. 5b, delivery of undiluted plasma at 24 hrs post infection (p.i.) demonstrated a partial protective effect. Finally, to assess the effects of CP treatment on SARS-CoV-2-induced anosmia, we treated infected mice with CP at 1 dpi and assessed mice for olfactory loss as described above (Fig. 4). Even though mice had minimal signs of clinical disease after CP treatment, by day 4 all mice exhibited profound anosmia (Fig. 4b, d, e).

**Fig. 5.**
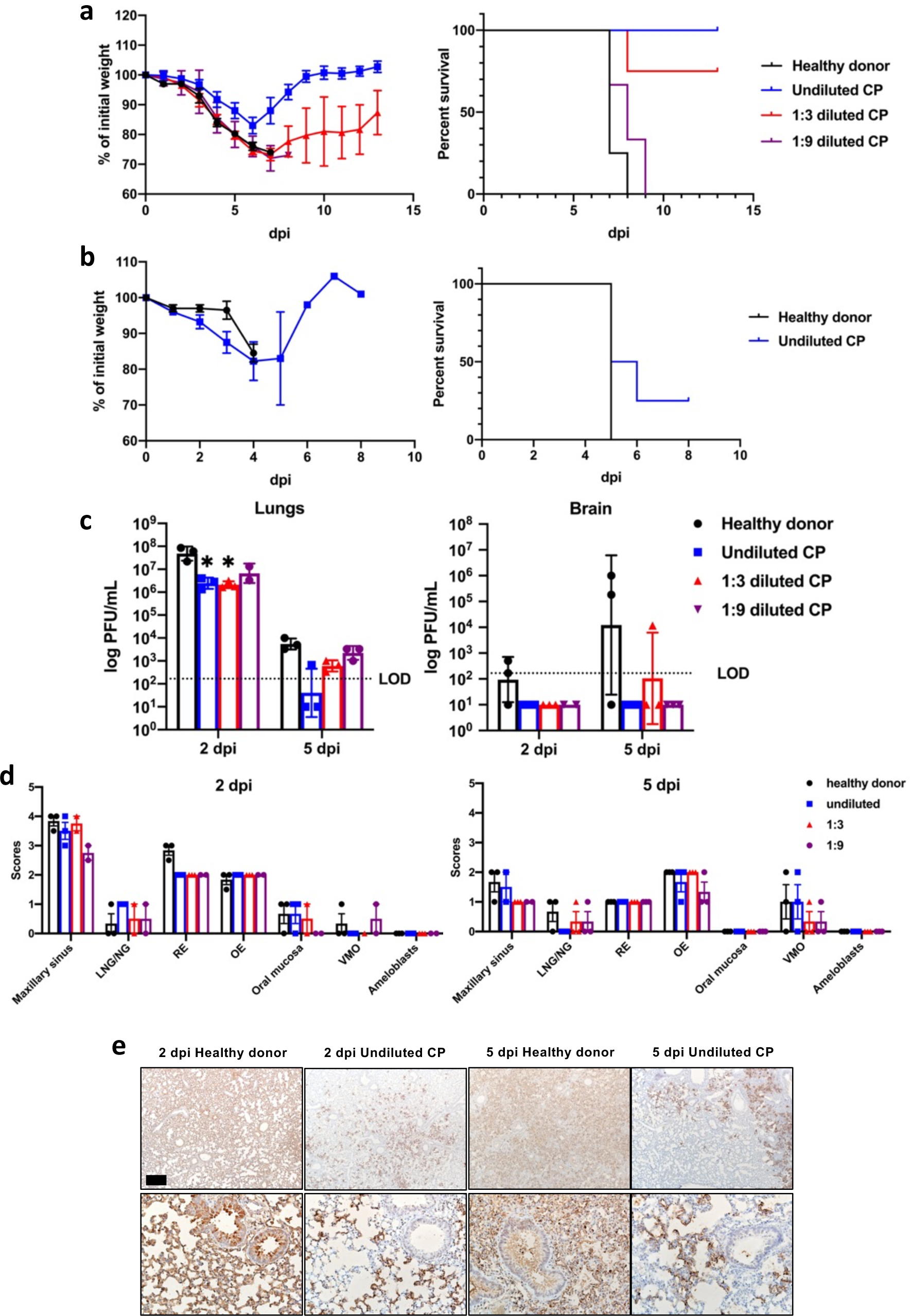
Effects of convalescent plasma on outcomes in SARS-CoV-2-infected K18-hACE2 mice. a. Percentage of initial weight (left panel) and survival (right panel) of infected K18-hACE2 mice receiving control serum (n=4, black), undiluted (n=4, blue), 1:3 diluted (n=4, red) and 1:9 diluted (n=3, purple) human convalescent plasma at 24 hours prior to challenge of 10^5^ PFU SARS-CoV-2. b. Percentage of initial weight (left panel), and survival (right panel) of K18-hACE2 mice receiving control plasma (n=2) or undiluted CP (n=4) at 24 hours after challenge with 10^5^ PFU SARS-CoV-2. c. Viral titers of CP-treated mice in the lungs (left panel) and brains (right panel) at 2 and 5 dpi. LOD=limit of detection. d. Scoring for N protein abundance in CP-treated mice in the nasal cavity at 2 (left panel) and 5 (right panel) dpi. 0 – none; 1 - rare <1%; 2 - multifocal or localized <33% cells; 3 - multifocal, coalescing, 33-66%; 4 - extensive >67%. e. N protein staining in the lungs of control or convalescent plasma treated mice at 2 and 5 days p.i. Bar = 370 µm (top) and 75 µm (bottom).

## DISCUSSION

When infected with SARS-CoV-2, K18-hACE2 mice developed a dose-dependent lung disease phenotype with features similar to severe human COVID-19. This includes diffuse alveolar damage, an influx of immune effector cells, tissue injury, lung vascular damage, and death. Remarkably, the mice also support SARS-CoV-2 replication in the sinonasal epithelium and associated with this pathology develop anosmia, a common feature of human disease. Furthermore, the uniformly fatal disease outcome with a 10^5^ inoculum was prevented by pre-treatment with CP from a COVID-19 patient. Notably, CP pre-treatment enhanced the kinetics of virus clearance but did not prevent initial infection of the lungs, damage to nasal respiratory and olfactory epithelia, or anosmia. SARS-CoV-2 infection of K18-hACE2 mice treated with CP or, potentially, neutralizing monoclonal antibodies will be especially useful for studies of anosmia because mice do not succumb to the infection, but like many infected patients with mild disease, have olfactory loss as a major manifestation^18^. Anosmia may result from damage to supporting sustentacular cells and not to olfactory sensory neurons, suggesting that the resulting inflammatory milieu, rather than direct neuronal damage is disease-causing. Therefore, we postulate that the observed anosmia has at least two possible explanations. First, the infection of sustentacular cells may disrupt signaling from olfactory sensory neurons (OSN) to the olfactory bulb. For example, sustentacular cells provide support to OSN by maintaining ion balance^28^, which may be altered either directly by sustentacular cell infection or indirectly via disorganization of the olfactory epithelium (OE), leading to loss of cilia from OSN and deficient signal transduction^29^. The second possibility is collateral damage to OSN resulting from pro-inflammatory cytokine secretion by SARS-CoV-2 infected sustentacular cells. These cells have been reported to secrete cytokines such as TNF-α, Il-1β, Il-1α, and CXCL2 in chronic rhinosinusitis and COVID-19 patients^30,31^. These are not mutually exclusive explanations.

Although both SARS-CoV and SARS-CoV-2 use ACE2 to gain entry into cells and both cause lung and brain disease, the disease manifestations are different. Most importantly, SARS-CoV caused a brain infection when as little as 3.2 PFU were administered intranasally^24^, while even 10^4^ PFU SARS-CoV-2 only variably infected the brain. A consequence of this difference in CNS susceptibility is that lung damage was obvious after SARS-CoV-2 infection, while it was obscured by the aspiration pneumonia that developed in SARS mice as a consequence of probable infection of the cardio-respiratory center in the medulla.

All K18-hACE2 mice succumbed to 10^5^ SARS-CoV-2, from lung and, in some cases, brain disease. This uniform lethality makes these mice useful for evaluation of anti-viral therapies and vaccines, and also sets a high standard for their efficacy. This is illustrated in Fig. 5, where mice treated with convalescent plasma were protected from lethal disease, but not infection. This absence of complete protection could reflect use of human convalescent plasma in mice, since some antibody effector functions are species-specific. Of note, while most mice receiving 1:3 or 1:9 dilutions of convalescent plasma had no evidence of brain infection (Fig. 5c), they succumbed to progressive lung disease. Infection with a range of virus inocula will be useful for evaluation of therapeutic interventions in a variety of pathological settings, as well as assessment of upper airway disease and anosmia. Therefore, K18-hACE2 mice, readily available from Jackson Laboratories, provide a useful model to study the pathogenesis of SARS-CoV-2-mediated disease and to evaluate interventions.

## SUPPLEMENTAL INFORMATION

is linked to the online version of the paper.

## ACKNOWLEDGEMENTS

This work is supported in part by grants from the National Institutes of Health USA (NIH) (P01 AI060699 (SP, PBM) and RO1 AI129269 (SP); and the Pathology Core, which is partially supported by the Center for Gene Therapy for Cystic Fibrosis (NIH Grant P30 DK-54759), and the Cystic Fibrosis Foundation. PBM is supported by the Roy J. Carver Charitable Trust.

## AUTHOR CONTRIBUTIONS

Conceptualization and writing – original draft, P.B.M., S.P.; Data curation, A.K.V.; Formal analysis, J.Z., L-Y R.W., K.L., M.E.O., A.K.V., D.K.M., P.B.M., and S.P.; Investigation, J.Z., L-Y R.W., K.L., M.E.O., C.W-L, A.K.V., C.M.K., D.K.M., P.B.M., and S.P.; Visualization, A.K.V, M.R.L., D.K.M.; Resources, D.K.M., P.B.M., and S.P.; Writing – review and editing, J.Z., L-Y R.W., S.P. and P.B.M.

## EXPERIMENTAL PROCEDURES

### Human subjects approval

Written informed consent was obtained from subjects to obtain plasma for participation in this study. The study was approved by the Institutional Review Board of the University of Iowa (IRB (#202003554 and #201402735).

### Mice, cells, and virus

Transgenic mice expressing human ACE2 were previously reported ^12^. Human ACE2 is expressed under control of the cytokeratin 18 promoter. The mice used in these studies were obtained from the Jackson Laboratory (034860-B6.Cg-Tg(K18-ACE2)2Prlman/J) and are congenic on the C57BL/6 background. Non-transgenic C57BL/6 mice were used as controls in some experiments.

All protocols were approved by the Institutional Animal Care and Use Committees of the University of Iowa. The 2019n-CoV/USA-WA1/2019 strain of SARS-CoV-2 (Accession number: MT985325.1) used in these studies was passaged on Calu-3 2B4 cells. Calu-3 2B4 cells were grown in MEM (GIBCO, Grand Island, NY) supplemented with 20% FBS. Vero E6 cells were grown in Dulbecco’s modified Eagle’s medium (DMEM, GIBCO, Grand Island, NY) supplemented with 10% fetal bovine serum (FBS).

### Infection of mice

Mice were lightly anesthetized with ketamine/xylazine and infected intranasally with the indicated amount of SARS-CoV-2 in a total volume of 50 µl DMEM. Animal weight and health were monitored daily. All experiments with SARS-CoV-2 were performed in a Biosafety Level 3 (BSL3) Laboratory at the University of Iowa.

### Nasal lavage

Mice were sedated with ketamine/xylazine (100 mg/kg ketamine/12.5 mg/kg xylazine). 100 µl of Carbachol (100 µg/ml) was administered I.P. and after approximately 15 minutes, nasal secretions were recovered. 20 µl secretions were recovered per mouse. Nasal secretions were immediately added to 500 µl Trizol, mixed and stored at −80°C until RNA was isolated.

### Virus titer by plaque assay

Virus or tissue homogenate supernatants were serially diluted in DMEM. 12 well plates of VeroE6 cells were inoculated at 37°C in 5% CO_2_ for 1 h and gently rocked every 15 min. After removing the inocula, plates were overlaid with 1.2% agarose containing 4% FBS. After 3 days, overlays were removed, and plaques visualized by staining with 0.1% crystal violet. Viral titers were quantified as PFU/mL tissue.

### Flow Cytometry

The following monoclonal antibodies were used: anti-mouse CD3e-BV421 (clone 145-2C11, Cat. No.: 562600), anti-mouse CD16/32 (clone 93, Cat. No.:101302), anti-mouse CD4-PercP (clone RM4-5, Cat. No.: 550954), anti-mouse CD8-APC-Cy7 (clone 53-6.7, Cat. No.: 100714), anti-mouse CD220-APC (clone RA3-6B2, Cat. No.: 553092), anti-mouse Ly6C-PerCP (clone HK1.4, Cat. No.: 128028), anti-mouse Ly6G-FITC (clone 1A8, Cat. No.: 127606), anti-mouse CD11b-BV510 (clone M1/70, Cat. No.: 101263), anti-mouse CD11c-BV421 (clone N418, Cat. No.: 117343), anti-mouse CD64-PE-Cy7 (X54-5/7.1), Biolegend, San Diego, CA; anti-mouse CD103-APC (clone 2E7, Cat. No.: 17-1031-80), anti-mouse TNF-FITC (clone MP6-XT22, Cat. No.: 11-7321-82), anti-mouse IFN-γ-APC (clone XMG1.2, Cat. No.: 25-7311-82), eBioscience, San Diego, CA; and mouse anti-C9 antibody (clone Rho 1D4, Cat. No.: MAB5356, EMD Millipore). For intracellular cytokine staining (ICS), lymphocytes were cultured in 96-well dishes at 37°C for 5-6 h in the presence of 2 μM peptide pool and brefeldin A (BD Biosciences). Cells were then labeled for cell-surface markers, fixed/permeabilized with Cytofix/Cytoperm Solution (BD Biosciences), and labeled with anti-IFN-γ and anti-TNF antibody. All flow cytometry data were acquired using a BD FACSVerse and analyzed with FlowJo software.

### Histology and Immunohistochemistry

Animals were anesthetized and perfused transcardially with PBS, followed by zinc formalin. Lungs were fixed in zinc formalin. For routine histology, tissue sections (∼4 µm each) were stained with hematoxylin and eosin. The following criteria were used for scoring edema, hyaline membrane formation and necrotic cellular debris: 0- none; 1- uncommon detection in <5% lung fields (200x); 2- detectable in up to 33% of lung fields; 3- detectable in up to 33-66% of lung fields; 4- detectable in >66% of lung fields. For scoring neutrophil infiltration: 0- within normal limits; 1-scattered PMNs sequestered in septa; 2- #1 plus solitary PMNs extravasated in airspaces; 3-#2 plus small aggregates in vessel and airspaces. For scoring mononuclear infiltrates, thrombosis and hemorrhage: 0-none; 1 - uncommon detection in <5% lung fields (200x); 2- detectable in up to 33% of lung fields; 3- detectable in up to 33-66% of lung fields; 4- detectable in >66% of lung fields.

For SARS-CoV-2 antigen detection, slides were incubated with blocking reagent (10% normal goat serum x 30 minutes) followed by rabbit monoclonal antibody against SARS-CoV2 N protein (1:20,000 dilution x 60 minutes, #40143-R019, Sino Biological US Inc., Wayne, PA, USA), then incubated with Rabbit Envision (Dako) and diaminobenzidine (Dako) as chromogen. Tissues were examined and scored in a post-examination method of masking by a boarded experimental pathologist ^32^. Ordinal scores for lesion parameters were assigned using the following tiers: 0 = within expected limits; 1 - uncommon, <5%; 2 - detectable in 5-33%; 3 - detectable in 34-66% and 4 - detectable in >66% of lung fields (200x objective magnification).

### Human convalescent plasma

High titer convalescent plasma was collected with subject consent under an IRB (#202003554) approved protocol that allows use of samples for research. The convalescent plasma donor was a 58 year old female who had molecularly confirmed COVID-19 more than 4 weeks prior to their donation. Following the donation, she tested positive for HLA antibodies so the plasma was not eligible for administration to patients and was diverted to research. Antibody testing (EUROIMMUN SARS-COV-2 ELISA (IgG)) performed on this donor was 9.8, well above the cutoff of 1.1 for a positive result. Neutralization titer using a luciferase-expressing SARS-CoV-2 S protein pseudovirus assay showed that the neutralization IC_50_ titer was 1:1,480. Control plasma was obtained from an expired plasma unit collected prior to COVID-19 spread in our area and this product was collected under an IRB (#201402735) approved protocol that allows for research use of these products.

### RNA isolation and qRT PCR

Total RNA was extracted from tissues (or nasal lavage) using Trizol (Invitrogen) according to the manufacturer’s protocol. Following a DNase treatment step 200 ng of total RNA was used as a template for first strand cDNA. The resulting cDNA was subjected to amplification of selected genes by real-time quantitative PCR using Power SYBR Green PCR Master Mix (Applied Biosystems). Average values from duplicates of each gene were used to calculate the relative abundance of transcripts normalized to HPRT and presented as 2^-ΔCT^. The primers used for cytokine and chemokines were previously reported^33^. For detection of viral genomes, the following primers were used to amplify the genomic RNA for the N protein: 2019-nCoV_N1-F: 5’-GAC CCC AAA ATC AGC GAA AT-3’; 2019-nCoV_N1-R: 5’-TCT GGT TAC TGC CAG TTG AAT CTG-3’. The following primers were used to amplify the subgenomic RNA for the E protein: F: 5’-CGATCTCTTGTAGATCTGTTCTC-3’; R: 5’-ATATTGCAGCAGTACGCACACA-3’.

### Pseudovirus neutralizing antibody assay

To determine neutralization activity of patient and mouse plasma, we used a luciferase reporter-based pseudovirus neutralization assay, which has a nonreplicative vesicular stomatitis virus backbone coated with the SARS-CoV-2 spike protein. Sera from SARS-CoV-2-infected K18-hACE2 mice were collected at the indicated time points and heat inactivated by incubation at 56°C for 30 min. Human convalescent plasma or mouse sera were serially diluted twofold in 96-well plates and incubated with the same volume of VSV pseudotyped with SARS-CoV-2 spike protein at 37°C for 1 h. Samples were placed onto Vero E6 cells and incubated at 37°C for 1 h to allow virus binding. After removal of the initial inoculum, cells were incubated for 24 hours. Neutralization was measured with a luciferase assay kit (Promega) and plotted relative to the value for control wells.

### Behavioral studies

#### Social Scent Discrimination Test

Social-scent discrimination task was designed to assess the capability of mice to discriminate social scents as described previously ^34^. Two identical tubes were separately sealed in a Ziploc bag in a large cage (used for housing rats (31 cm x 26 cm x 22 cm)) containing bedding from female mice and from the experimental cage overnight. Next, mice were released in a fresh cage containing the tubes placed at two different corners. Sniffing latency (sniffing and exploring the tubes) was calculated within a 5 min period. One trial on each mouse was performed each day with the position of tubes changed daily. Data were recorded in the BSL3 facility from a distance of approximately 1 meter.

#### Buried Food Test

Evaluation of odor detection was carried out by a buried food test (bft) as described previously^34^. Briefly, food was confirmed prior to initiation of the experiments. Mice were sensitized to the food for 10-15 minutes on the day of the experiment. The food pellet was then buried approximately 1 cm below the surface of bedding in a fresh cage. One trial per mouse per day was performed with a changed position of food on a daily basis. The latency to locate and uncover the buried food pellet was recorded with a stopwatch. Mice were allowed to explore the cage for 4 min and if they could not locate the food, the time was recorded as 4 min. The experiment was repeated daily.

### Statistical analysis

ANOVA and Student’s t tests were used to analyze differences in mean values between groups using GraphPad Prism 7. All results are expressed as mean ± standard error (SEM) and were corrected for multiple comparisons. For behavioral studies, data from the social scent discrimination test, and buried food test were analyzed by two-way ANOVA and one-way ANOVA tests, respectively. *P* values of <0.05 were considered statistically significant. (*\*, P values of* ≤*0*.*05. **, P values of* ≤*0*.*005. ***, P values of* ≤*0*.*0005. ****, P values of* ≤*0*.*0001)*.

### Data availability

The data supporting the findings of this study are documented within the paper and are available from the corresponding authors upon request.

## FIGURE LEGENDS

**Extended Data Fig. 1. Histological analysis of extrapulmonary tissue in SARS-CoV-2-infected K18-hACE2 mice**. Mice were sacrificed at days 0, 4 and 6 p.i. and tissues prepared for histological examination (n=3/4 per group). Liver (a), spleen (b), kidney (c), small intestine (d), and colon (e) were studied. Pathological changes were minor and only observed in the liver. In the liver, all mice had some blood vessels filled with clear space or aggregates variably composed of erythrocytes / platelets (insets). Rare vessels had evidence of eosinophilic fibrillar material consistent with fibrin thrombi (arrows, inset in middle panel) adherent along the vascular wall.

**Extended Data Fig. 2**. Gating strategy for identification of immune cells in lungs is shown.

## REFERENCES

1 Zhou, P. et al. A pneumonia outbreak associated with a new coronavirus of probable bat origin. Nature 579, 270–273, doi:10.1038/s41586-020-2012-7 (2020).

2 Chan, J. F. et al. Simulation of the clinical and pathological manifestations of Coronavirus Disease 2019 (COVID-19) in golden Syrian hamster model: implications for disease pathogenesis and transmissibility. Clin Infect Dis, doi:10.1093/cid/ciaa325 (2020).

3 Munster, V. J. et al. Respiratory disease and virus shedding in rhesus macaques inoculated with SARS-CoV-2. bioRxiv, doi:10.1101/2020.03.21.001628 (2020).

4 Rockx, B. et al. Comparative pathogenesis of COVID-19, MERS, and SARS in a nonhuman primate model. Science 368, 1012–1015, doi:10.1126/science.abb7314 (2020).

5 Shi, J. et al. Susceptibility of ferrets, cats, dogs, and other domesticated animals to SARS-coronavirus 2. Science 368, 1016–1020, doi:10.1126/science.abb7015 (2020).

6 Sia, S. F. et al. Pathogenesis and transmission of SARS-CoV-2 in golden hamsters. Nature 583, 834–838, doi:10.1038/s41586-020-2342-5 (2020).

7 Wan, Y., Shang, J., Graham, R., Baric, R. S. & Li, F. Receptor Recognition by the Novel Coronavirus from Wuhan: an Analysis Based on Decade-Long Structural Studies of SARS Coronavirus. J Virol 94, doi:10.1128/JVI.00127-20 (2020).

8 Hassan, A. O. et al. A SARS-CoV-2 Infection Model in Mice Demonstrates Protection by Neutralizing Antibodies. Cell, doi:10.1016/j.cell.2020.06.011 (2020).

9 Sun, J. et al. Generation of a Broadly Useful Model for COVID-19 Pathogenesis, Vaccination, and Treatment. Cell, doi:10.1016/j.cell.2020.06.010 (2020).

10 Dinnon, K. H. et al. A mouse-adapted SARS-CoV-2 model for the evaluation of COVID-19 medical countermeasures. bioRxiv, doi:10.1101/2020.05.06.081497 (2020).

11 Sun, S. H. et al. A Mouse Model of SARS-CoV-2 Infection and Pathogenesis. Cell Host Microbe 28, 124–133 e124, doi:10.1016/j.chom.2020.05.020 (2020).

12 McCray, P. B., Jr. et al. Lethal infection in K18-hACE2 mice infected with SARS-CoV. J Virol 81, 813–821 (2006).

13 Tseng, C. T. et al. SARS coronavirus infection of mice transgenic for the human angiotensin-converting enzyme 2 (hACE2) virus receptor. J Virol 81, 1162–1173 (2006).

14 Yang, X. H. et al. Mice transgenic for human angiotensin-converting enzyme 2 provide a model for SARS coronavirus infection. Comp Med 57, 450–459 (2007).

15 Chow, Y. H. et al. Development of an epithelium-specific expression cassette with human DNA regulatory elements for transgene expression in lung airways. Proc Natl Acad Sci U S A 94, 14695–14700 (1997).

16 Ackermann, M. et al. Pulmonary Vascular Endothelialitis, Thrombosis, and Angiogenesis in Covid-19. N Engl J Med 383, 120–128, doi:10.1056/NEJMoa2015432 (2020).

17 Feldstein, L. R. et al. Multisystem Inflammatory Syndrome in U.S. Children and Adolescents. N Engl J Med 383, 334–346, doi:10.1056/NEJMoa2021680 (2020).

18 Spinato, G. et al. Alterations in Smell or Taste in Mildly Symptomatic Outpatients With SARS-CoV-2 Infection. JAMA, doi:10.1001/jama.2020.6771 (2020).

19 Chow, Y. H. et al. Targeting transgene expression to airway epithelia and submucosal glands, prominent sites of human CFTR expression. Mol Ther 2, 359–367, doi:10.1006/mthe.2000.0135 (2000).

20 Bradley, B. T. et al. Histopathology and ultrastructural findings of fatal COVID-19 infections in Washington State: a case series. Lancet 396, 320–332, doi:10.1016/S0140-6736(20)31305-2 (2020).

21 Channappanavar, R. et al. Dysregulated type I interferon and inflammatory monocyte-macrophage responses cause lethal pneumonia in SARS-CoV-infected mice. Cell Host Microbe 19, 181–193, doi:10.1016/j.chom.2016.01.007 (2016).

22 Mathew, D. et al. Deep immune profiling of COVID-19 patients reveals distinct immunotypes with therapeutic implications. Science, doi:10.1126/science.abc8511 (2020).

23 Poyiadji, N. et al. COVID-19-associated Acute Hemorrhagic Necrotizing Encephalopathy: Imaging Features. Radiology 296, E119–E120, doi:10.1148/radiol.2020201187 (2020).

24 Netland, J., Meyerholz, D. K., Moore, S., Cassell, M. & Perlman, S. Severe acute respiratory syndrome coronavirus infection causes neuronal death in the absence of encephalitis in mice transgenic for human ACE2. J Virol 82, 7264–7275 (2008).

25 Bilinska, K., Jakubowska, P., Von Bartheld, C. S. & Butowt, R. Expression of the SARS-CoV-2 Entry Proteins, ACE2 and TMPRSS2, in Cells of the Olfactory Epithelium: Identification of Cell Types and Trends with Age. ACS Chem Neurosci 11, 1555–1562, doi:10.1021/acschemneuro.0c00210 (2020).

26 Joyner, M. J. et al. Early safety indicators of COVID-19 convalescent plasma in 5,000 patients. J Clin Invest, doi:10.1172/JCI140200 (2020).

27 Casadevall, A., Joyner, M. J. & Pirofski, L.A. SARS-CoV-2 viral load and antibody responses: the case for convalescent plasma therapy. J Clin Invest, doi:10.1172/JCI139760 (2020).

28 Vogalis, F., Hegg, C. C. & Lucero, M. T. Ionic conductances in sustentacular cells of the mouse olfactory epithelium. J Physiol 562, 785–799, doi:10.1113/jphysiol.2004.079228 (2005).

29 Bryche, B. et al. Massive transient damage of the olfactory epithelium associated with infection of sustentacular cells by SARS-CoV-2 in golden Syrian hamsters. Brain Behav Immun, doi:10.1016/j.bbi.2020.06.032 (2020).

30 Imamura, F. & Hasegawa-Ishii, S. Environmental Toxicants-Induced Immune Responses in the Olfactory Mucosa. Front Immunol 7, 475, doi:10.3389/fimmu.2016.00475 (2016).

31 Torabi, A. et al. Proinflammatory Cytokines in the Olfactory Mucosa Result in COVID-19 Induced Anosmia. ACS Chem Neurosci 11, 1909–1913, doi:10.1021/acschemneuro.0c00249 (2020).

32 Meyerholz, D. K. & Beck, A. P. Fundamental Concepts for Semiquantitative Tissue Scoring in Translational Research. ILAR J 59, 13–17, doi:10.1093/ilar/ily025 (2018).

33 Li, K. et al. Middle East Respiratory Syndrome Coronavirus Causes Multiple Organ Damage and Lethal Disease in Mice Transgenic for Human Dipeptidyl Peptidase 4. J Infect Dis 213, 712–722, doi:10.1093/infdis/jiv499 (2016).

34 Yang, M. & Crawley, J. N. Simple behavioral assessment of mouse olfaction. Curr Protoc Neurosci Chapter 8, Unit 8 24, doi:10.1002/0471142301.ns0824s48 (2009).

