## Supplemental figure 1, Supplemental figure 2, Table 1 for "K18-hACE2 Mice for Studies of COVID-19 Treatments and Pathogenesis Including Anosmia"

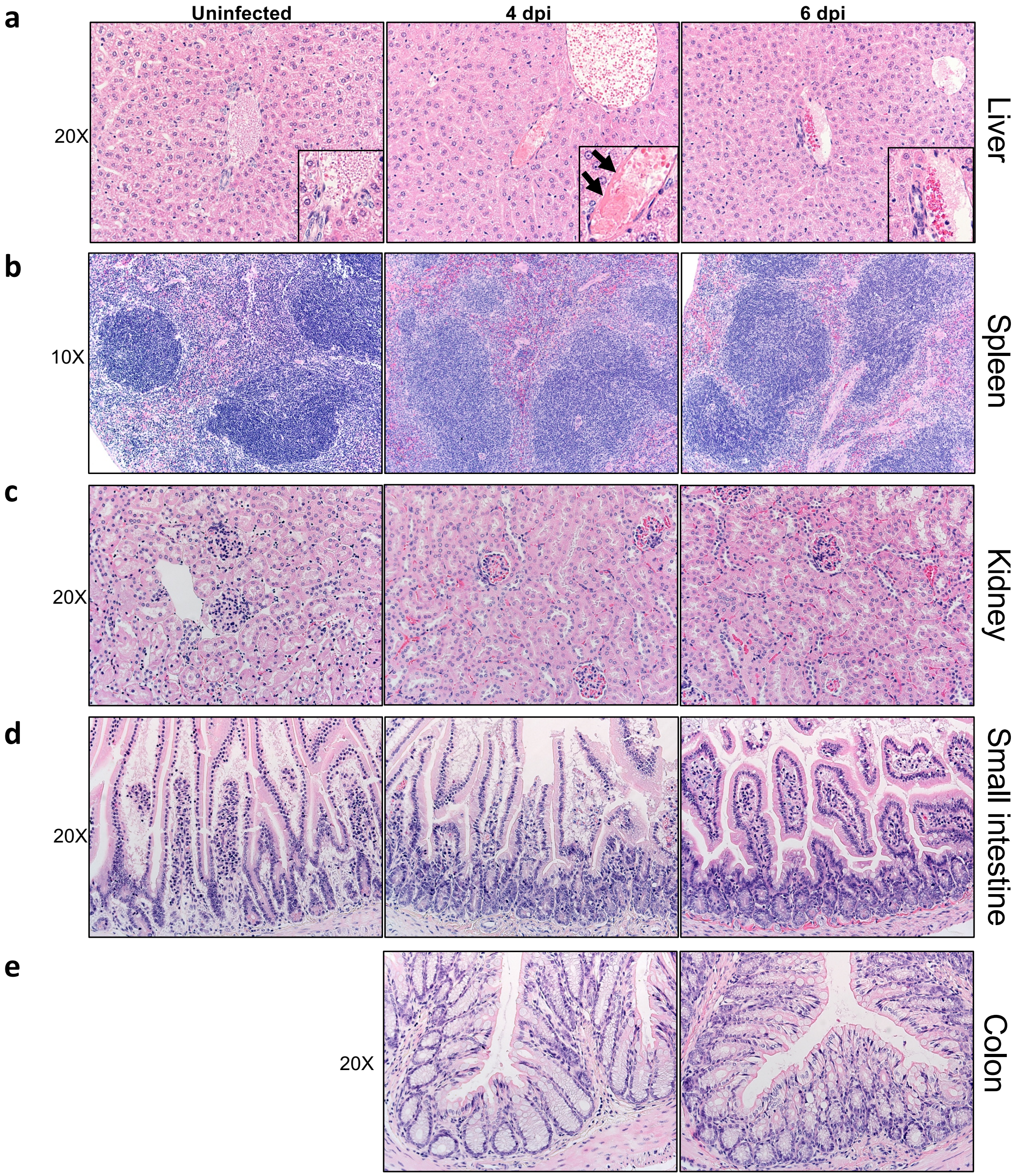

**Extended Data Fig. 1. Histological analysis of extrapulmonary tissue in SARS-CoV-2-infected K18-hACE2 mice.** Mice were sacrificed at days 0, 4 and 6 p.i. and tissues prepared for histological examination (n=3/4 per group). Liver (a), spleen (b), kidney (c), small intestine (d), and colon (e) were studied. Pathological changes were minor and only observed in the liver. In the liver, all mice had some blood vessels filled with clear space or aggregates variably composed of erythrocytes / platelets (insets). Rare vessels had evidence of eosinophilic fibrillar material consistent with fibrin thrombi (arrows, inset in middle panel) adherent along the vascular wall.

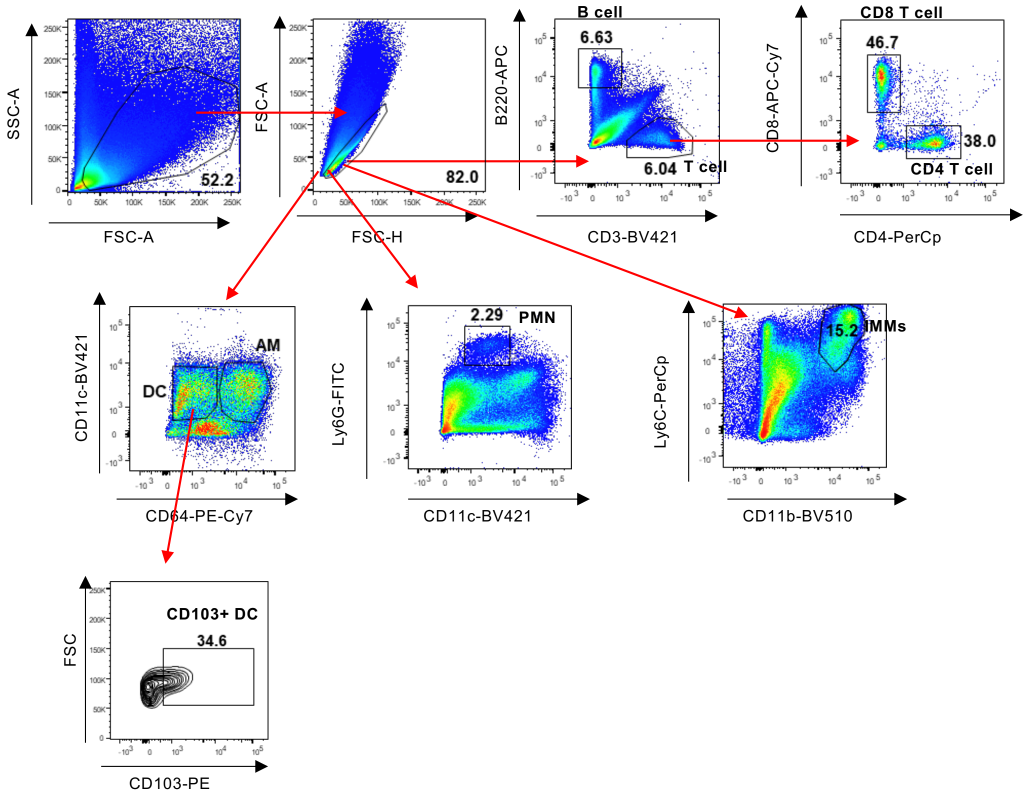

**Extended Data Fig. 2**. Gating strategy for identification of immune cells in lungs is shown.

**Extended Figure Table 1. Distribution of SARS-CoV-2 antigen in brain**

### Olfactory Bulb

#### Main Olfactory Bulb

| | |
|---|---|
| Glomerular Layer | - |
| Granule Layer | - |
| Inner Plexiform Layer | - |
| Mitral Layer | + |
| Outer Plexiform Layer | - |

#### Accessory Olfactory Bulb

| | |
|---|---|
| Glomerular Layer | - |
| Granular Layer | - |
| Mitral Layer | ++ |
| AON Dorsal | +++ |
| AON Medial | ++ |
| AON lateral | ++ |
| Taenia Tecta | + |

### Thalamus

| | |
|---|---|
| Ventral Posterolateral Nucleus | +++ |
| Ventral Posteromedial Nucleus | +++ |
| Ventral Medial Nucleus | ++ |
| Ventral Anterior-Lateral Complex | +++ |
| Anterior group of the Dorsal Thalamus | +++ |
| Medial group of the Dorsal Thalamus | +++ |
| Midline group of the Dorsal Thalamus | +++ |
| Reticular Nucleus | +++ |
| Geniculate Group | +++ |

### Hypothalamus

| | |
|---|---|
| Supraoptic Nucleus | +++ |
| Paraventricular Nucleus | +++ |
| Arcuate Nucleus | +++ |
| Suprachiasmatic Nucleus | +++ |
| Subfornical Organ | +++ |
| Anterior Nucleus | +++ |
| Premammillary Nucleus | +++ |

### Cerebral Cortex

| | |
|---|---|
| Frontal | +++ |
| Infralimbic | ++ |
| Prelimbic | ++ |
| Lateral Orbital | ++ |
| Cingulate | ++ |
| Parietal | +++ |
| Retrosplenial | ++ |
| Temporal | +++ |
| Occipital | ++ |
| Perirhinal | ++ |
| Piriform | +++ |

### Basal Ganglia

| | |
|---|---|
| Caudate Putamen | +++ |
| Globus Pallidus | +++ |
| Ventral Pallidum | +++ |
| Lateral Preoptic | +++ |
| Nucleus Accumbens | +++ |
| Fundus Striati | +++ |
| Olfactory Tubercle | +++ |

### Midbrain

| | |
|---|---|
| Superior Colliculus | ++ |
| Inferior Colliculus | ++ |
| Nucleus Brachium | ++ |
| Nucleus Sagulum | ++ |
| Para Bigeminal Nucleus | ++ |
| Midbrain Trigeminal Nucleus | ++ |
| Substantia Nigra | +++ |
| Ventral Tegmental Area | ++ |
| Reticular Nucleus | +++ |
| Cuneiform Nucleus | ++ |
| Red Nucleus | ++ |
| Oculomotor Nucleus III | ++ |
| Edinger-Westphal Nucleus | ++ |
| Trochlear Nucleus IV | ++ |
| Ventral Tegmental Nucleus | ++ |
| Pedunculopontine Nucleus | ++ |

### Hind Brain

#### Pons

| | |
|---|---|
| Nucleus of Lateral Lemniscus | ++ |
| Trigeminal Nucleus | ++ |
| Parabrachial Nucleus | ++ |
| Dorsal Tegmental Nucleus | ++ |
| Pontine Nuclei | ++ |

| | |
|---|---|
| **Hippocampus** | (+) |
| **Cerebellum** | - |

| | |
|---|---|
| Supratrigeminal Nucleus | ++ |
| Superior Central Nucleus Raphe | ++ |
| Locus Ceruleus | ++ |
| Laterodorsal Tegmental Nucleus | ++ |
| <u>Medulla</u> | |
| Area Postrema | +++ |
| Cochlear Nuclei | ++ |
| Dorsal Column Nuclei | ++ |
| External Cuneate Nucleus | + |
| Nucleus Ambiguus | ++ |
| Solitary Tract Nucleus | ++ |
| Lateral Reticular Nucleus | ++ |
| Magnocellular Reticular Nucleus | ++ |
| Medullary Reticular Nucleus | ++ |
| Parvicellular Reticular Nucleus | ++ |
| Nucleus X | ++ |
| Hypoglossal Nucleus XII | ++ |

4 brains were analyzed at 6dpi.

(+) Represents presence of antigen in some samples only.

Neuroanatomical reference : ALLEN Brain Atlas

http://atlas.brain-map.org/atlas?atlas=2#atlas=2&plate=100883770&structure=168&x=7797.5&y=4023.9999997615814&zoom=-3&resolution=10.47&z=5

AON- Anterior Olfactory Nucleus

+ <++<+++ represents relative density of antigen presence; - No antigen detected
